# Customized feminine hygiene wash containing postbiotics from *Lactobacillus* spp. to treat Urinary Tract Infections (UTI)

**DOI:** 10.1101/2025.01.24.634779

**Authors:** Veena G Nair, David Raj Chellappan, Remya Devi Durai, Y.B.R.D Rajesh, Dhiviya Narbhavi, A Anupriya, N Prabhusaran, N Saisubramanian

**Author notes:** Corresponding Author: Dr. Sai Subramanian N, Antimicrobial Resistance Lab, Centre for Research in Infectious diseases, School of Chemical and Biotechnology, SASTRA Deemed to be University, Thanjavur 613401, Tamil Nadu, India. **Email:**.

## Abstract

The rising prevalence of antimicrobial resistance has intensified the search for innovative therapeutic strategies, particularly in the prevention of urinary tract infections (UTIs). This study presents the development and evaluation of a novel postbiotics vaginal wash formulated to prevent UTIs by utilizing metabolites derived from indigenous vaginal Lactobacillus spp. The primary objective was to create a cost-effective, stable, and non-invasive solution targeting uropathogenic bacteria. Key metabolites, including tryptamine, (-)-terpinen-4-ol, and itaconic anhydride, were identified from cell free supernatant of vaginal *Lactobacillus* and incorporated into a poloxamer 407-based formulation. *In vitro* assays demonstrated significant bioactivity against uropathogenic bacteria, effectively inhibiting bacterial colonization and biofilm formation. Preclinical validation was conducted using BALB/c mice models to assess both the safety and efficacy of the vaginal wash. Results indicated a substantial reduction in infection rates among treated mice, with no observed adverse effects, confirming the formulation’s safety profile. In conclusion, this novel postbiotics vaginal wash represents a promising non-invasive therapeutic approach for UTI prevention. By harnessing bioactive metabolites from human vaginal Lactobacillus spp., the formulation offers a potential solution to combat antimicrobial resistance while improving women’s health outcomes. Further clinical studies are warranted to validate these findings and explore broader applications in clinical practice, paving the way for new strategies in managing UTIs and enhancing overall female health.

## Introduction

The growing issue of antibiotic resistance on a global scale is a pressing concern, significantly compromising the effectiveness of commonly used antibiotics against widespread bacterial infections. The 2022 Global Antimicrobial Resistance and Use Surveillance System (GLASS) report, also identifies extremely high levels of resistance in common pathogens. The global rapid situation assessment based on median and updated country-level reported rates shows that in 76 countries the projected resistance burden on third generation cephalosporins is as high as 42% rate of resistance in *E. coli*, and for methicillin-resistant *Staphylococcus aureus* it is up to 35%. Particularly for UTIs linked to *E. coli*, disturbingly one in five cases were less susceptible to conventional antibiotics such as ampicillin, co-trimoxazole and fluoroquinolones in 2020(1). This worrisome trend is making it increasingly challenging to effectively combat and treat common infections, emphasizing the urgent need for global strategies to address antibiotic resistance.

Over the recent years, there has been a noticeable surge in the pharmaceutical industry’s interest in developing new formulations that incorporate beneficial microorganisms, often tailored to the specific needs of the host.(2) Nevertheless, the heightened focus on the clinical use of probiotics necessitates consideration when administering them to populations with health challenges. Recently, the Food and Drug Administration created a new category called “live biotherapeutic products” (LBP). As a result, the documentation and demonstration of quality, safety and efficacy for new products (including LBPs) are adapted to specific properties such as risk profiles stemming from both target population characteristics along with those related more directly to product strains or components. This approach enables a comprehensive assessment of the global benefit-risk ratio concerning their intended use.(3) The pharmacological aspects of probiotics are more intricate compared to conventional drugs due to their multifaceted impact on host cells, mucosal organs, and the entire organism. Evaluating their effects is challenging as they operate through diverse mechanisms, either individually or synergistically(4). The reports describe different ways vaginal lactobacilli can exert their functions as follows: (a) producing antimicrobial substances such as organic acids, hydrogen peroxide, bacteriocins and biosurfactants; or enzymes like arginine deaminase; (b) adhesion to epithelial cells, mucus components or extracellular matrix for colonization and domination of this ecological niche environment by autoaggregation co-aggregations, biofilms in the vagina mucosa, competitive exclusion competition for nutrients and immunomodulation.(5, 6)

Although probiotics are generally considered safe, concerns persist about the potential risks associated with administering live bacteria in specific situations, such as during pregnancy or while preparing for motherhood(7, 8). Additionally, ensuring the stability of probiotic products—so that they maintain enough viable bacteria during handling, transportation, and storage—presents practical challenges in industrial production and marketing(9). As a result of these challenges, there has been considerable research focus on postbiotics, inactivated probiotics, and the products formed during the breakdown of probiotic bacteria. This research is increasingly demonstrating their potential to promote health and treat diseases(10). Postbiotics were defined as “a preparation of inanimate microorganisms and/or their components that confers a health benefit on the host” by The International Scientific Association for Probiotics and Prebiotics (ISAPP) published its consensus statement regarding Postscript in 2021 (9). In recent years, there has been increasing research focus on the significant effects of *Lactobacillus* bioactive compounds (postbiotics) on vaginal health (10–13). One of the most important biological activities for postbiotics is to reduce vaginal pH to 3.8–4, which has antimicrobial effects resulting in significant inhibition of pathogenic microorganisms(14–16). Research indicates that vaginal *Lactobacilli* and the bioactive compounds they produce, including hydrogen peroxide, lactic acid, surfactants, and ribosomally synthesized antimicrobial peptides (bacteriocins), are critical in suppressing the growth and proliferation of vaginal pathogens(17, 18). Most likely due to bacteriocins, cell-free supernatants exerted the highest scale of antibacterial activity (19). For instance, Cell free supernatants from *Lactobacillus* and *Bifidobacterium* genera have demonstrated antibacterial properties against enteroinvasive *E. coli*(20). Exopolysaccharides (EPS) from *Bifidobacterium bifidum* are used to promote the growth of *Lactobacilli* and other anaerobes & inhibit *Enterococci*, *Enterobacteriacea*, and *B. fragilis* (21). Furthermore, the antibacterial metabolite reuterin, produced by *Lactobacillus reuteri*, is believed to act by oxidizing thiol groups in pathogenic gut bacteria(22).

While most studies on postbiotics highlight common metabolites such as lactic acid, our earlier works have identified unique bioactive compounds from human vaginal *Lactobacillus* with distinct therapeutic properties. These metabolites include tryptamine, a biogenic amine(23); (-)-terpinen-4-ol, a naturally occurring monoterpene (Manuscript under review); and itaconic anhydride, the cyclic form of itaconic acid (Manuscript under Review). Unlike typical postbiotics, these compounds exhibit specific bioactivities such as antibiofilm, efflux inhibitory, and antipersister activities.

Tryptamine disrupts the exopolysaccharide matrix of bacterial biofilms, enhancing their susceptibility to antimicrobial agents. (-)-Terpinen-4-ol demonstrates significant efflux inhibitory activity, counteracting the mechanisms employed by uropathogenic bacteria to expel antibiotics. Itaconic anhydride works synergistically with (-)-terpinen-4-ol to mitigate the virulence of persister cells in uropathogenic *E. coli*, providing a robust defense against persistent bacterial infections. Leveraging these findings, we have developed a customized vaginal wash formulation incorporating these bioactive postbiotics with a poloxamer 407 base, a biocompatible polymer known for its stability and suitability for mucosal applications. Preclinical validation conducted on BALB/c mice models has demonstrated the efficacy and safety of this innovative formulation, underscoring its potential for clinical translation. This novel formulation represents a significant advancement in UTI prevention, addressing the urgent need for new therapies that can combat antimicrobial resistance and improve women’s health.

## Materials and Methods

### *In vitro* cell line infection and toxicity study

T24 bladder epithelial cells and RAW macrophages (NCCS, Pune were utilized to assess the efficacy and cytotoxicity of various treatments targeting bacterial infections. T24 cells, derived from transitional bladder carcinoma, were chosen for their close resemblance to primary human bladder epithelial cells, making them an ideal model for studying urinary tract infections (UTIs). The cells were grown in McCoy’s 5A medium, enhanced with 10% fetal bovine serum (FBS), and plated in a 24-well plate at a density of 1×10^5^ cells/mL. To provide optimal conditions for growth, the T24 cells were incubated for 16 hours at 37°C in a 5% CO2/95% air atmosphere.

Following a 2-hour incubation period at 37°C with 5% CO2, the bacterial culture was prepared at a multiplicity of infection (MOI) of one and added to 24-well plates. Treatments were administered for 24 hours, after which cells were rinsed with sterile PBS to eliminate unbound bacteria. Cells were then trypsinized to detach and recover adhered bacteria, and bacterial counts were determined via serial dilution on Luria-Bertani agar plates. Each assay was performed in triplicate and repeated twice for validation.

For cytotoxicity assessments on RAW macrophages, the MTT assay was used. Cells grown to 70–80% confluence were trypsinized, resuspended in fresh media, and seeded into 96-well plates. Following 24 hours of incubation, fresh media containing treatment compounds and MTT (0.5 mg/mL) was replaced. The formazan crystals were dissolved with DMSO after the two-hour incubation period, and the cell viability was evaluated by measuring the absorbance at 595 nm to assess cell viability(24).

### Biofilm formation

A 96-well microtiter plate was used, with 200 μL of test medium and 20 μL of a 48-hour LB broth culture as the inoculum. After incubating at 37°C for 48 hours under static conditions, unbound cells were washed with sterile PBS. Biofilm-bound cells were stained with 1% Crystal Violet (CV) for 15 minutes, washed with distilled water, and the bound dye was extracted with 70% ethanol. The absorbance of the extracted CV was measured at 595 nm. Viable cells were determined by scraping the biofilms from the wells, performing a 10-fold dilution, and spread plating to enumerate colony-forming units (CFU). (25) Treatments were administered with a triple combination of postbiotics at different pH levels.

### Rheological analysis

The viscosity of the formulation was measured using a Brookfield Viscometer (LVDV-II + Pro) with Spindle No. 64 at different temperatures. The formulation was equilibrated at the initial temperature and its viscosity was measured. The sample was then subjected to incremental temperature increases, with viscosity readings taken at each step after allowing the formulation to stabilize. The results, indicating viscosity changes with temperature, demonstrated the formulation’s suitability for prolonged retention and enhanced therapeutic efficacy, as a significant increase in viscosity with rising temperature suggests better performance under therapeutic conditions.

### *In vivo* infection study using BALB/c Mice

The reported animal experiments are confirmed to be compliant with ARRIVE guidelines by the Institutional Animal Ethical Committee of SASTRA Deemed to be University (Approval number: (740/SASTRA/IAEC/RPP)).

The *In vivo* study was organized into five groups (N=6 per group): disease control, placebo, customized vaginal wash, probiotic wash, and commercial wash. A pilot study was conducted before the main study to optimize the experimental parameters and determine the appropriate study duration. The main study, spanning 14 days, involved vaginal induction of *E. coli* UTI89 GFP in BALB/c mice by administering 20 µL of a bacterial suspension (10^6^CFU/mL) directly into the vaginal canal using a sterile pipette under aseptic conditions, ensuring precise and uniform colonization. Following induction, the mice were monitored for signs of infection, such as inflammation and vaginal discharge, with body weight recorded regularly. Treatment with the vaginal wash formulations was applied daily. Urine samples were collected on days 8, 11, and 14, and vaginal discharge samples were collected on days 3, 4, 5, 6, 8, 10, and 12. Blood samples were collected for serum creatinine assays (AGAPPE Diagnostics Ltd, Mumbai, India) to assess renal function. Mice were sacrificed, and their organs were homogenized to enumerate bacterial load.

For the preparation of the probiotic wash, 0.01 M lactic acid/sodium lactate buffer (pH 3.2) was cooled to 4 °C. To this buffer, 18% Poloxamer and freeze-dried bacterial powder were slowly added with continuous agitation. *Lactobacillus jensenii* was introduced at a concentration 10^9^ CFU/mL, and the mixture was left at 4 °C until a clear solution was obtained, achieving a final pH of 3.8.

The Poloxamer-407 formulation containing postbiotics was prepared by combining 18% Poloxamer with tryptamine (4 µg/mL), (-)-Terpinen-4-ol (5 µg/mL), and itaconic anhydride (8 µg/mL). For the commercial vaginal wash, 1 mL of the product was diluted with 99 mL of distilled water to create a solution suitable for administration in BALB/c mice. This dilution was performed to ensure safety and minimize potential irritation, as well as to adhere to the volume constraints appropriate for the species used in this study

## Results

### Triple combination of postbiotics reduces intracellular bioburden of *E. coli* UTI89 in T24 bladder cells, inhibits biofilm *invitro* and exhibits non-toxicity in Raw macrophages

The study evaluated the efficacy of postbiotic metabolites—tryptamine of 4 µg/ml, (-)-terpinen-4-ol (5 µg/ml), and itaconic anhydride (8 µg/ml) —in reducing bacterial load using an *In vitro* infection in T24 bladder cells and preformed biofilm assays (Figure.1A). These concentrations were selected based on our previous research, which indicated that these specific levels of tryptamine, (-)-terpinen-4-ol, and itaconic anhydride exhibited optimal antimicrobial activity against relevant bacterial strains. This prior work demonstrated that these concentrations effectively reduced bacterial load while maintaining cell viability, thereby supporting their use in the current study. Treatments included individual metabolites, synergistic combinations, and a triple-compound mixture. Notably, the combination of all three metabolites resulted in a significant 9-log reduction in bacterial load, underscoring their potent synergistic effect in combating bacterial infections, particularly within T24 bladder cells (Figure.1B). Additionally, in vitro toxicity studies using Raw macrophages were conducted to assess the safety of these compounds. The MTT assay revealed no significant cytotoxicity at concentrations of 4 µg/ml for tryptamine, 5 µg/ml for (-)-terpinen-4-ol, and 10 µg/ml for itaconic anhydride, indicating that these compounds are safe within the tested concentration range (Supplementary Figure.1). These findings suggest that the triple combination of postbiotics not only effectively reduces bacterial burden but also maintains a safety profile, making it a promising candidate for therapeutic applications.

**Figure 1.**
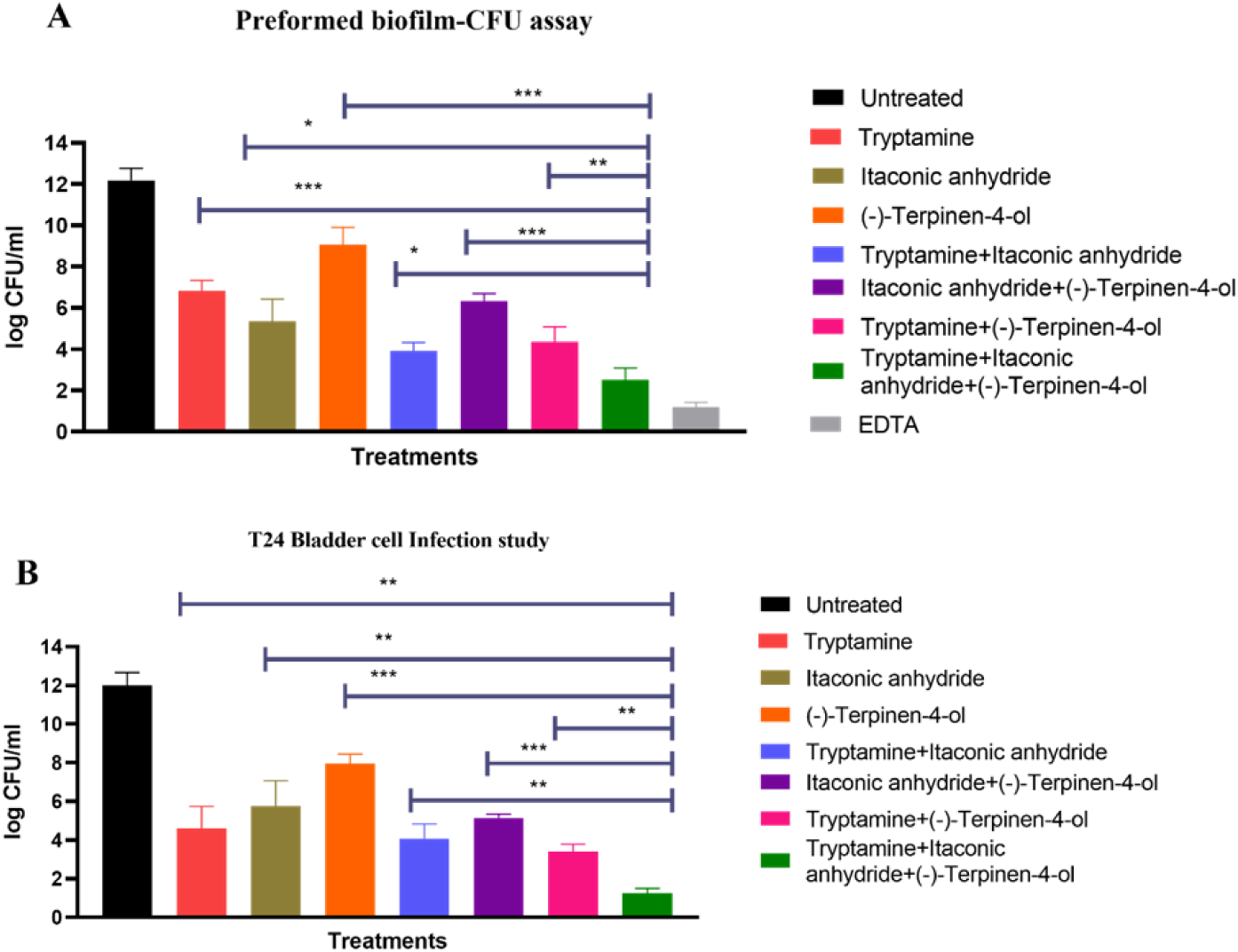
*In vitro* cell line infection study with T24 Bladder cells of the metabolites and its synergistic effects. A) Biofilm CFU Reduction in Preformed Biofilms Treated with Individual and Combined Metabolites B) Synergistic Effects of Metabolite Combinations on Bacterial Load in T24 Bladder Cells. One tail t test was performed to determine the significance, **p<0.01, ***p<0.001, n=3, EDTA (0.1%) used as a positive control for preformed biofilm.

### Development of a customized vaginal wash for enhanced intravaginal delivery of vaginal *Lactobacillus* metabolites

Poloxamer 407 (procured from Sigma-Aldrich), a copolymer known for its unique liquid-to-gel transition upon temperature alteration, forms the base of the formulation (Figure. 3). Poloxamer 407, has a molecular weight of approximately 12.6 kDa and is hydrophilic in nature owing to its polyoxyethylene content (∼70%) it serves as the key component of the formulation. This copolymer’s thermosensitive behavior, transitioning from a liquid to a gel state in response to temperature changes, enables precise control over gelation kinetics. The addition of Tryptamine (4 µg/ml), (-)-terpinen-4-ol (5 µg/ml), and itaconic anhydride (8µg/ml) further enhances the formulation’s therapeutic efficacy.

The optimal concentration of poloxamer for vaginal drug delivery systems is typically in the range of 18-21% w/w. At an optimized concentration of 18%, Poloxamer 407 confers the formulation with ideal rheological properties, ensuring optimal spreadability and retention within the vaginal region. At room temperature (25 °C), the formulation exhibits low viscosity, facilitating ease of application and mucosal coverage (Figure. 2B). Upon exposure to body temperature (37 °C), the formulation undergoes in situ gelation, ensuring prolonged retention within the vaginal environment (Figure. 2A). The rheological study using Brookfield Viscometer (LVDV-II + Pro, Brookfield Engineering, USA with spindle no. 64) demonstrated that the viscosity of the formulation increases significantly with the rise in temperature, indicating its suitability for prolonged retention and enhanced therapeutic efficacy (Figure 2C).

**Figure 2.**
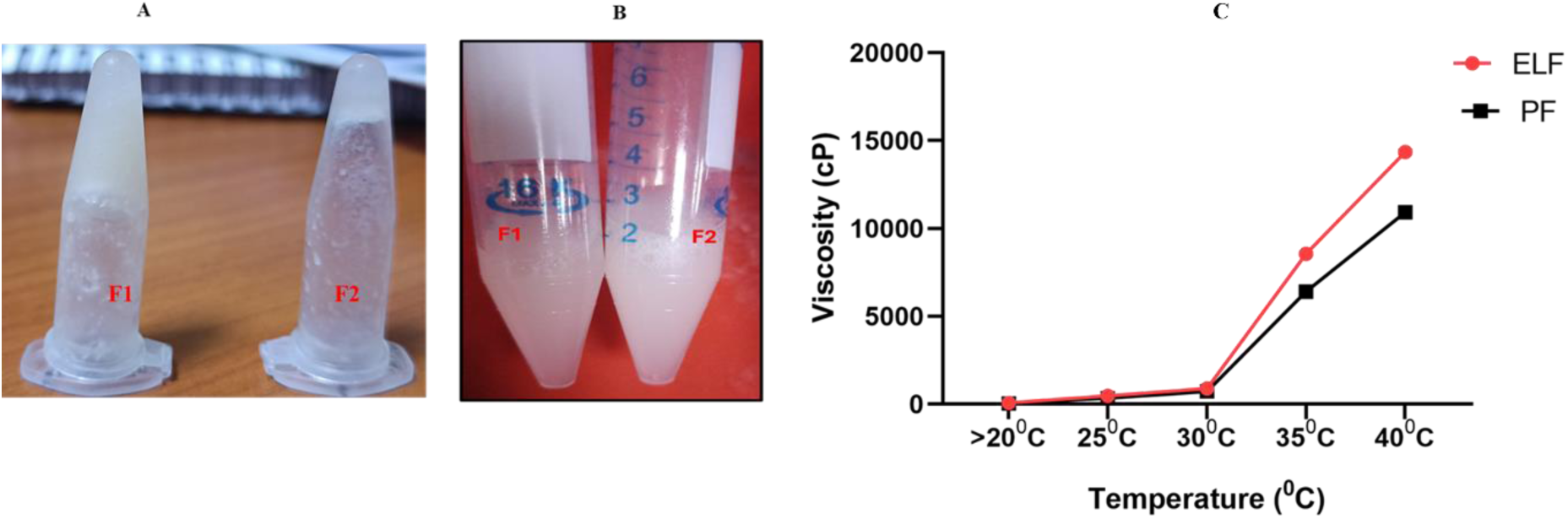
Sol-Gel Transformation of the customized vaginal wash formulation using Poloxamer-407, where F1 is the *Lactobacillus* extracted metabolites incorporated in 18% Poloxamer 407 and F2 is the procured metabolites incorporated in 18% Poloxamer 407 A) Gel form at 37^0^C B) Solution form at below 20 ^0^ C C) Rheological analysis of the formulation to check the viscosity using Brookfield Viscometer (LVDV-II + Pro, Brookfield Engineering, USA) with spindle no. 64. ELF means Extracted metabolites from *Lactobacillus* incorporated in poloxamer 407 and PLF procured compounds incorporated in poloxamer 407.

### Biofilm inhibition and bacterial growth suppression by the Customized Vaginal Wash

The findings from our study demonstrate the effectiveness of the customized vaginal wash formulation in significantly reducing biofilm formation and bacterial viability across a range of pH levels (Figure. 3A), highlighting its robust inhibitory capabilities. The observed decrease in biofilm density and structural integrity, as evidenced by microscopy (Figure.3B) and crystal violet assays (Supplementary Figure3C-D), suggests that the formulation effectively disrupts the biofilm matrix, a key factor in preventing persistent bacterial colonization and infection. The 9-log reduction in colony-forming units (CFUs) (Figure 3E) further confirms the formulation’s capacity to lower bacterial viability within the biofilm, suggesting that it not only inhibits biofilm matrix formation but also actively reduces the existing bacterial population (Figure 3E).

**Figure. 3.**
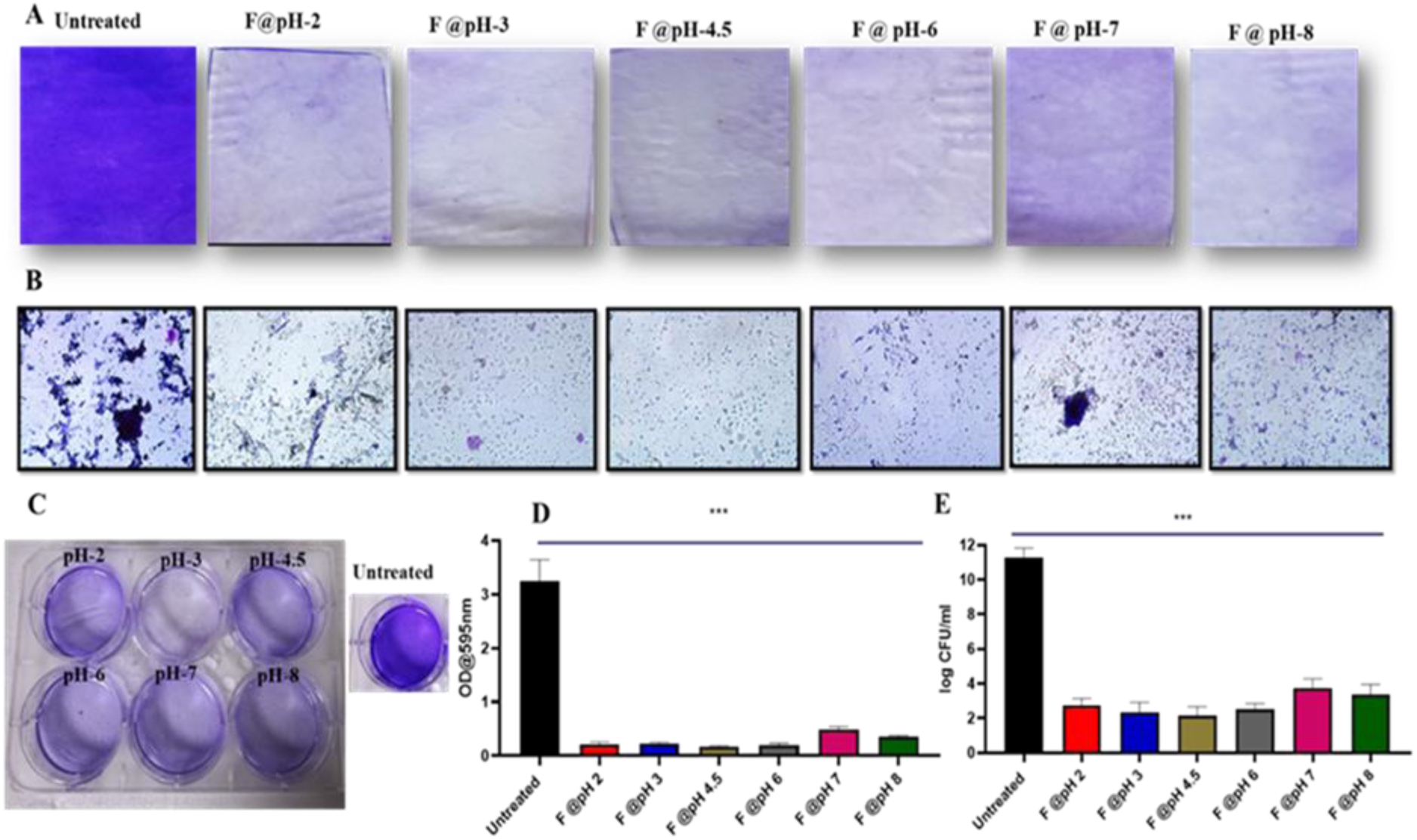
Inhibition of Biofilm formation matrix on cover glass by the customized vaginal wash formulation. A) Biofilm formation after treatment with different pH (2, 3, 4.5, 6, 7 and 8) of customized formulation B) Microscopic images of biofilm after treatment with different pH of customized formulation C) Plate showing Crystal violet assay for biofilm D) Quantification of crystal violet assay E) Colony forming units. One tail *t test* was performed to determine the significance., ***p<0.001, n=3

Complementing these findings, scanning electron microscopy (SEM) and fluorescence imaging were employed to assess bacterial growth inhibition following treatment with the customized formulation. SEM images revealed a significant reduction in bacterial density, coupled with noticeable morphological abnormalities in the treated groups compared to the untreated control (Figure. 4A). Fluorescence imaging further demonstrated decreased bacterial fluorescence intensity, indicating diminished bacterial viability post-treatment (Figure. 4B).

**Figure. 4.**
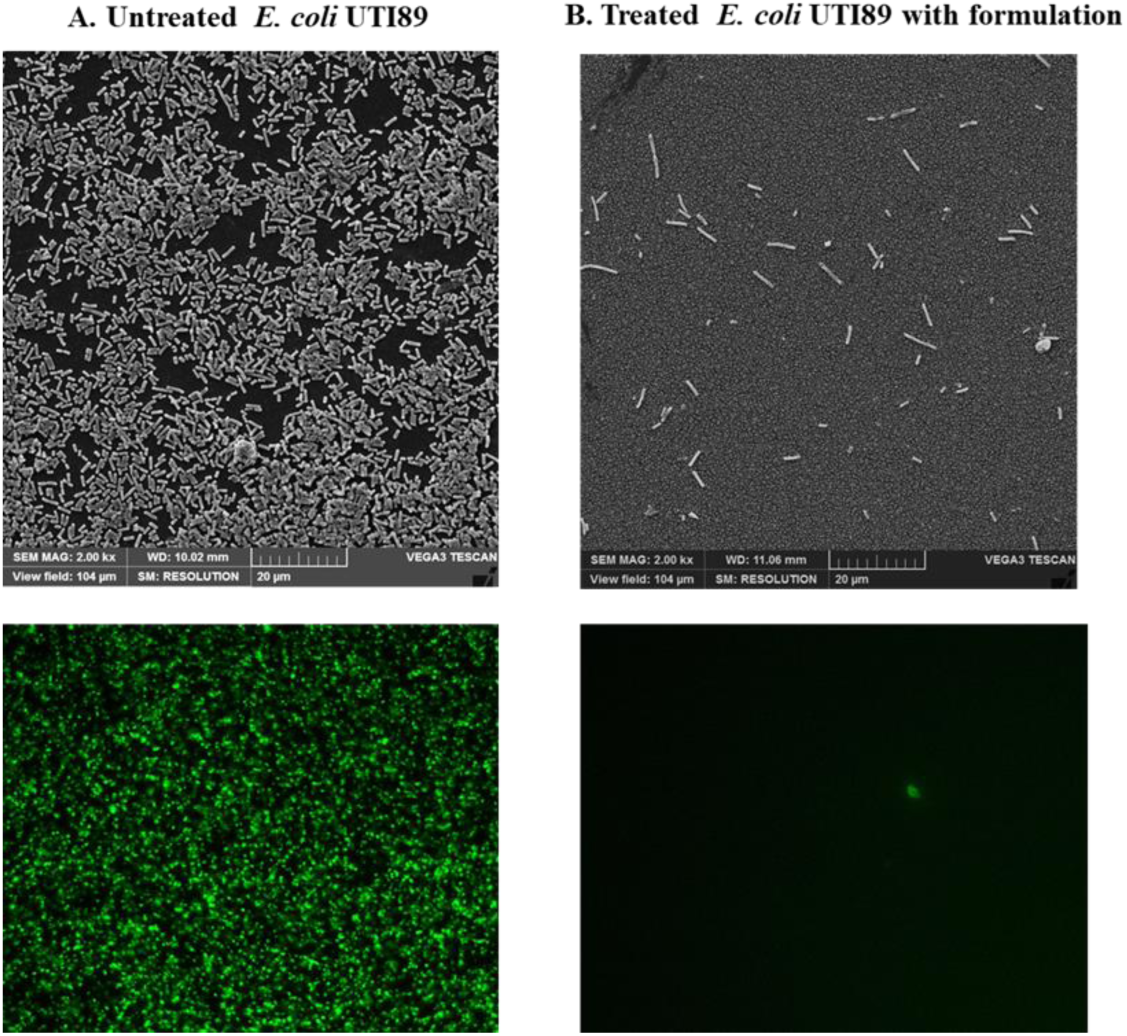
Scanning Electron Microscope and Fluorescent images. shows the inhibition of bacterial growth after treating with the formulation. A. Untreated *E. coli* UTI89, B. Treated with formulation. Scale bar =20µm

### Customized vaginal wash formulation retains its stability and biological activity for 3 months at 4°C

Results revealed that despite the prolonged storage period at 4°C for 3 months, the customized *Lactobacillus*-derived wash retained its ability to prevent biofilm formation of clinically relevant bacterial strains, including *E. coli* UTI89, *E. coli* CFT073, *E. coli* F11, *Klebsiella pneumoniae*, Methicillin-Resistant *Staphylococcus aureus* (MRSA), and *Pseudomonas aeruginosa*. (Supplementary Figure 2A) and there was ∼ 9 log reduction in the growth of all tested bacterial strains (Supplementary Figure 2B). This broad-spectrum efficacy underscores the wash’s potential utility in combating various infections encountered in clinical settings.

### Customized Vaginal Wash prevents vaginal inflammation in BALB/c mice without affecting body weight

In this study, the efficacy of various vaginal wash formulations was evaluated using a BALB/c mouse model of vaginal infection. The study included five groups (N=6 per group): disease control, placebo, customized vaginal wash, probiotic wash, and commercial wash.

Detailed schematics of the animal study are provided in Figure 5. On day 0, a 20 µL suspension containing 10^6^ CFU/mL of *E. coli* UTI89 GFP was instilled into the vaginal cavity of BALB/c mice to induce infection. The inoculation was performed under aseptic conditions to ensure precise bacterial colonization within the vaginal canal. Treatment with the respective vaginal wash formulations began on day 3 post-infection, continuing through day 14 (Supplementary Figure. 3).

**Figure 5:**
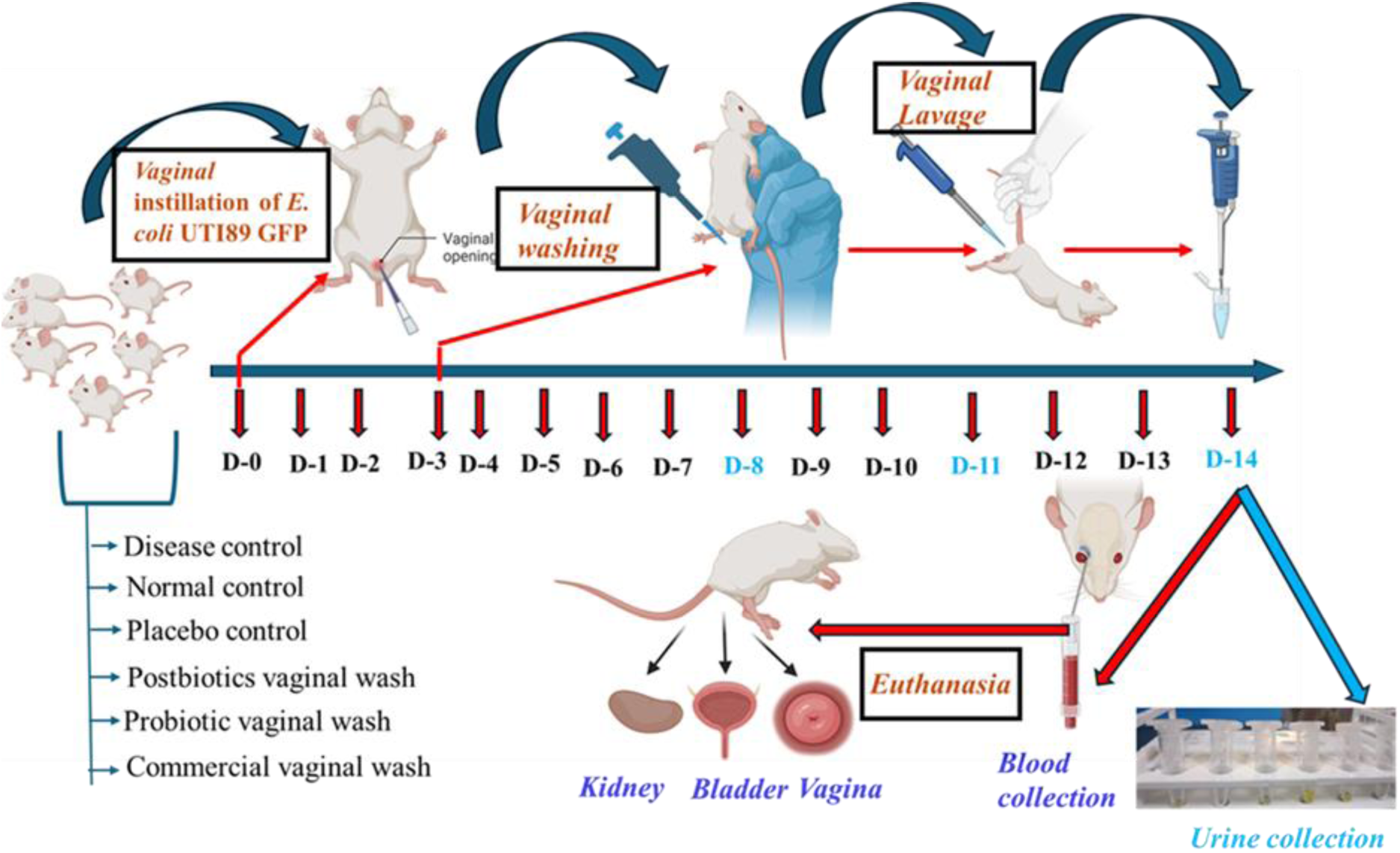
Schematics of Animal Study

By day 8, observations revealed significant differences among the treatment groups. The disease control, probiotic wash, and commercial wash groups exhibited pronounced inflammation, including redness, swelling, and the formation of papules. In contrast, the customized vaginal wash-treated group showed significantly reduced inflammation, with vaginal health closely resembling that of the normal control group (Figure.6A).

Throughout the study, the body weight of the mice was monitored. While the baseline weight ranged between 35-40 grams, by day 8, the disease control, probiotic wash, and commercial wash groups exhibited a substantial weight loss of approximately 10 grams. Notably, mice treated with the customized vaginal wash maintained their baseline weight, indicating better overall health and the efficacy of the customized formulation in managing the infection (Figure.6B).

**Figure 6.**
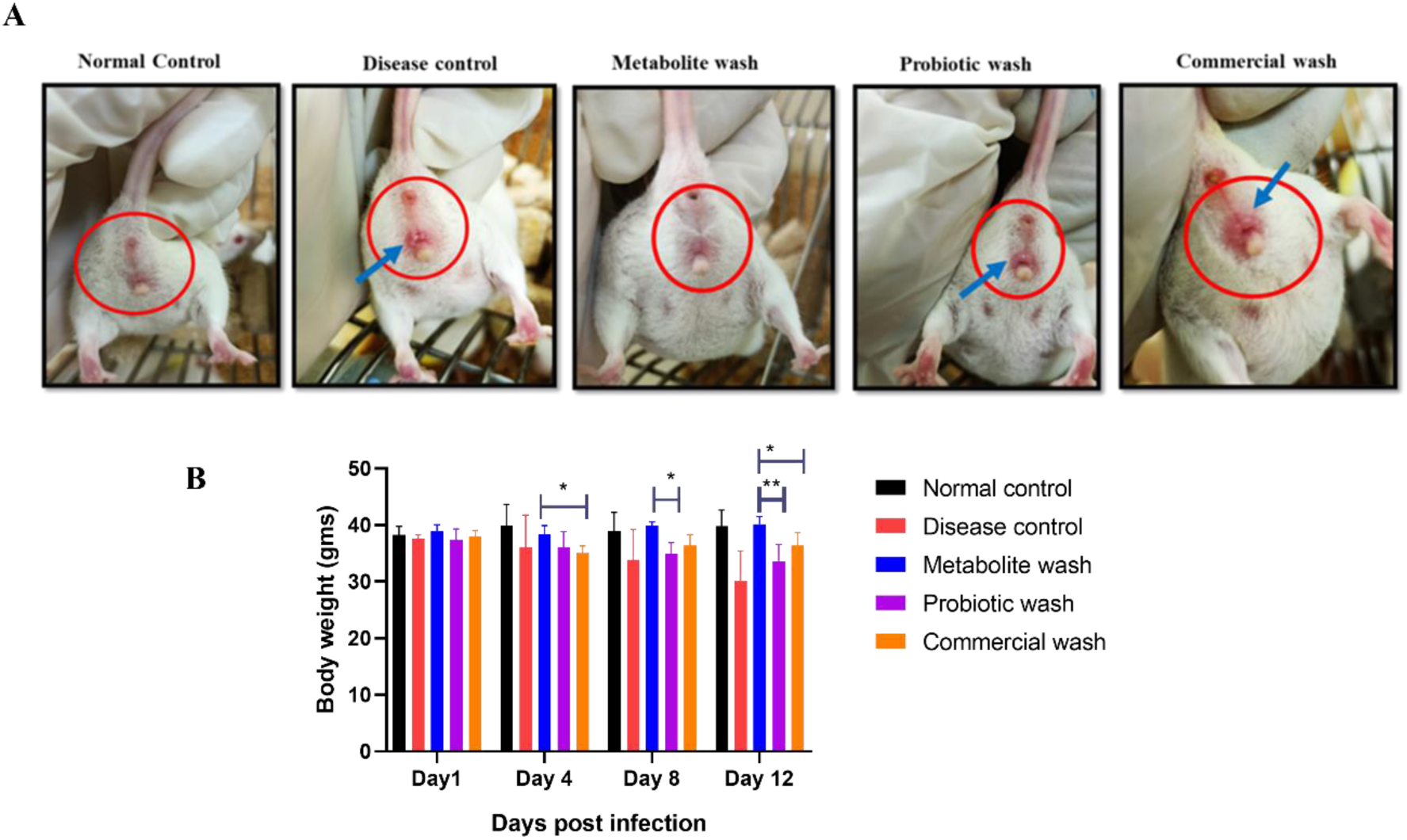
**Effect of vaginal wash formulations on inflammation and body weight in BALB/c Mice infected with *E. coli* UTI89 GFP**. A) Inflammation of vagina of BALB/c Mice, Blue arrow mark indicated the inflammation B) Body weight measurements of BALB/c mice. One tail t test was performed to determine the significance. *p<0.05, **p<0.01. Data represented as a mean ± standard error of the mean (SEM) (N= 6).

### Customized vaginal wash reduced bacterial load in vaginal discharge

Following formulation treatment, bacterial load in the vaginal discharge of BALB/c mice was quantified by plate counts on Day 14 post infection (Figure 7A), Since *E. coli* UTI89 GFP strain was used for infection, it was directly employed for fluorescent imaging (Figure 7B). Plate counts (Figure 7A) revealed that the metabolite-based formulation achieved a significant 8-10 log reduction in bacterial count compared to the untreated control, probiotic formulation, and commercial wash, demonstrating its efficacy in inhibiting bacterial growth and colonization. Both the probiotic wash and commercial wash displayed ∼ 2 log reduction in cell counts relative to untreated infected control. Similarly, fluorescent images on Day 6, Day 8, and Day 12 post-infection revealed that, relative to commercial wash, which failed to curtail the growth of *E. coli* UTI89 strain, metabolite wash completely mitigated *E. coli* UTI89 growth (Figure 7B).

**Figure. 7.**
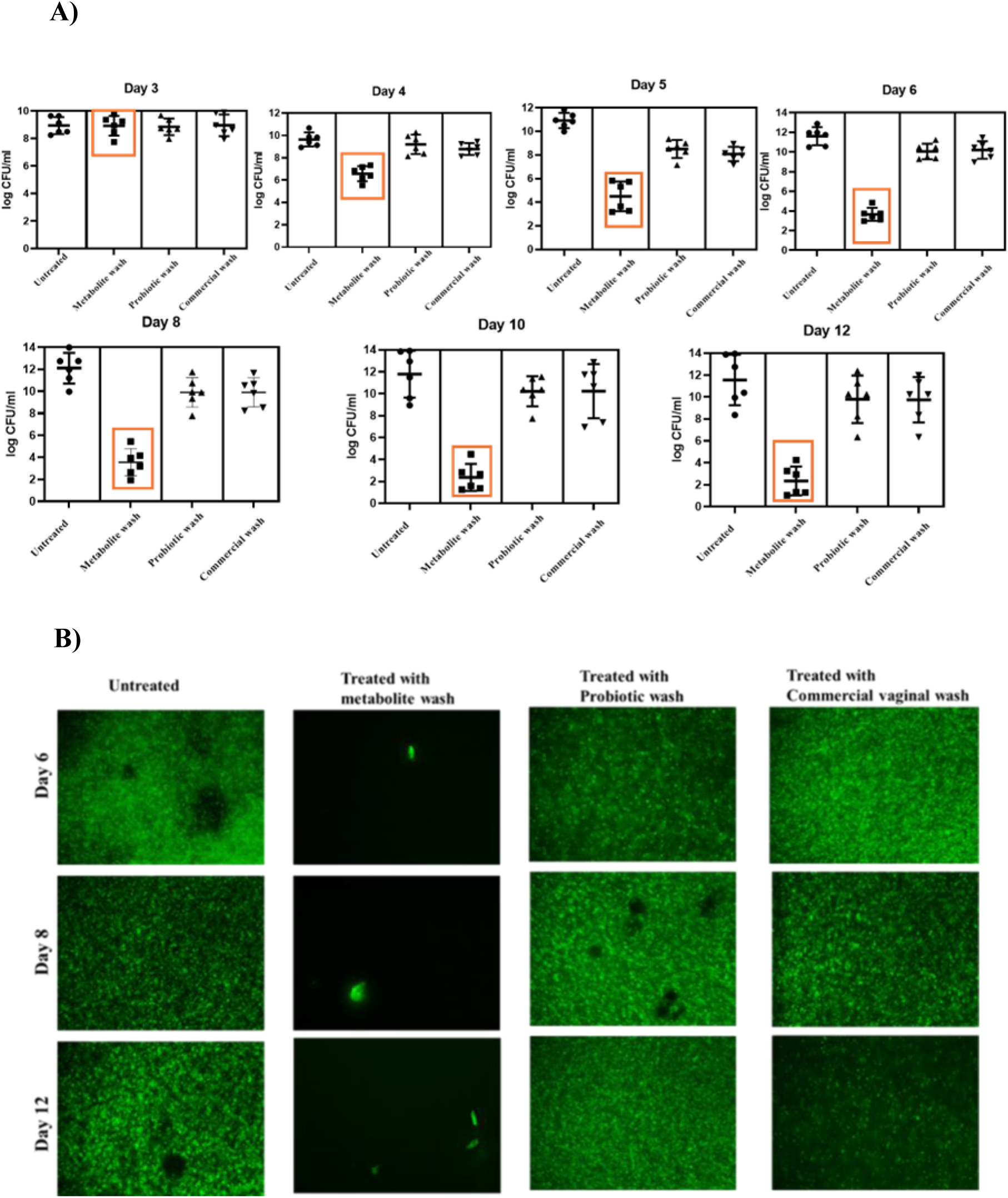
**Bacterial loads in vaginal discharge of BALB/c Mice post infection**. A) Quantifications were performed in triplicate and represented as a mean ± standard error of the mean (SEM) (N= 6) B) Fluorescent images showing the bacterial load in the vaginal discharge of BALB/c Mice at Day 4, Day 8, and Day 12 post infection. (N= 6)

### Absence of bacteria in urine/ organs after treatment with customized vaginal wash

We assessed the efficacy of a metabolite-based wash in preventing urinary tract infections (UTIs) and related complications in BALB/c mice. Infection was induced using *E. coli* UTI89 GFP, and bacterial load in urine was quantified at multiple time points post-infection (Figure. 82A). Fluorescent imaging of urine samples collected on Day 8, Day 11, and Day 14 revealed bacterial colonization dynamics within the urinary tract (Figure. 8B). Remarkably, no *E. coli* UTI89 GFP was detected in the urine of mice treated with the metabolite wash, indicating successful prevention of UTI development.

**Figure. 8.**
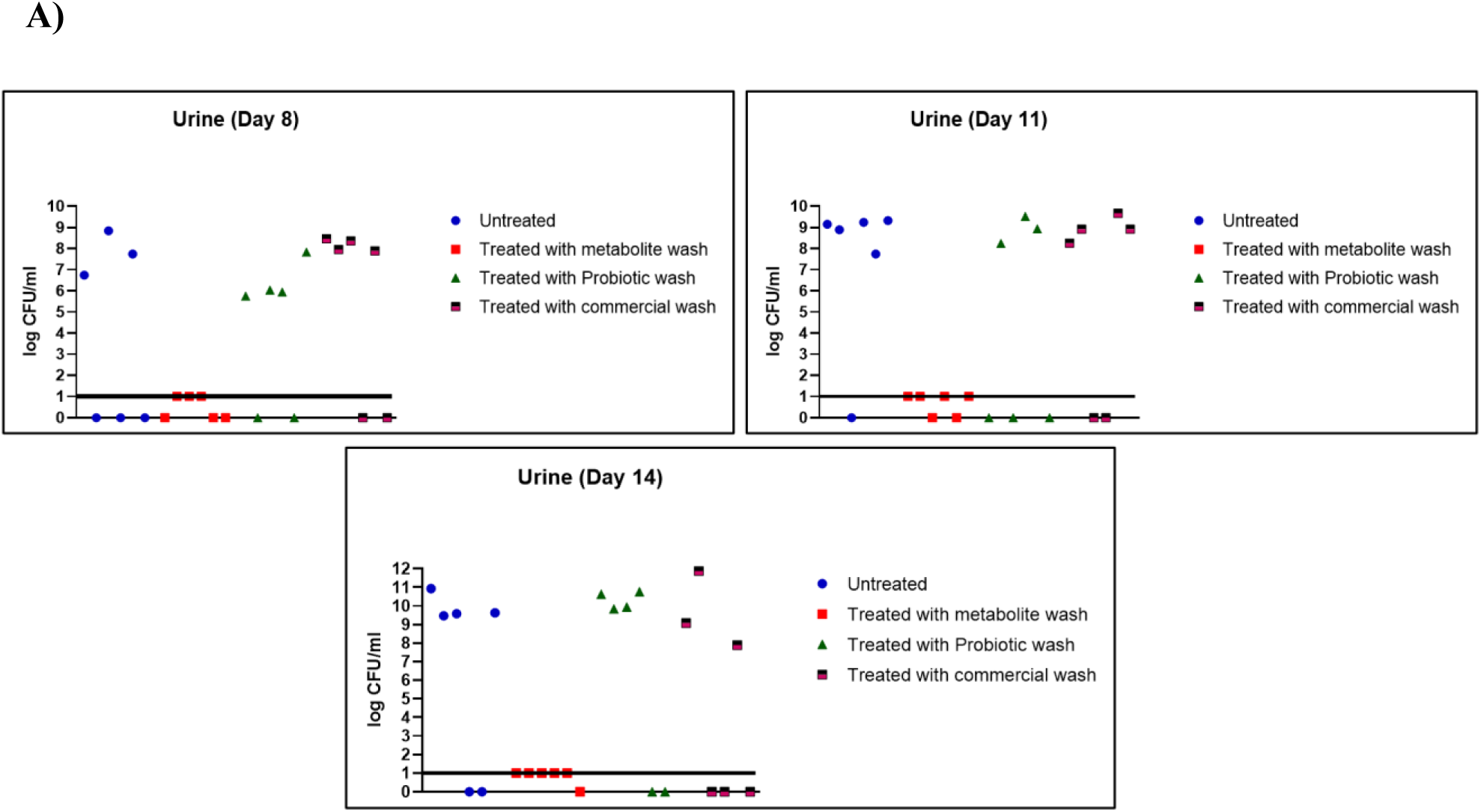
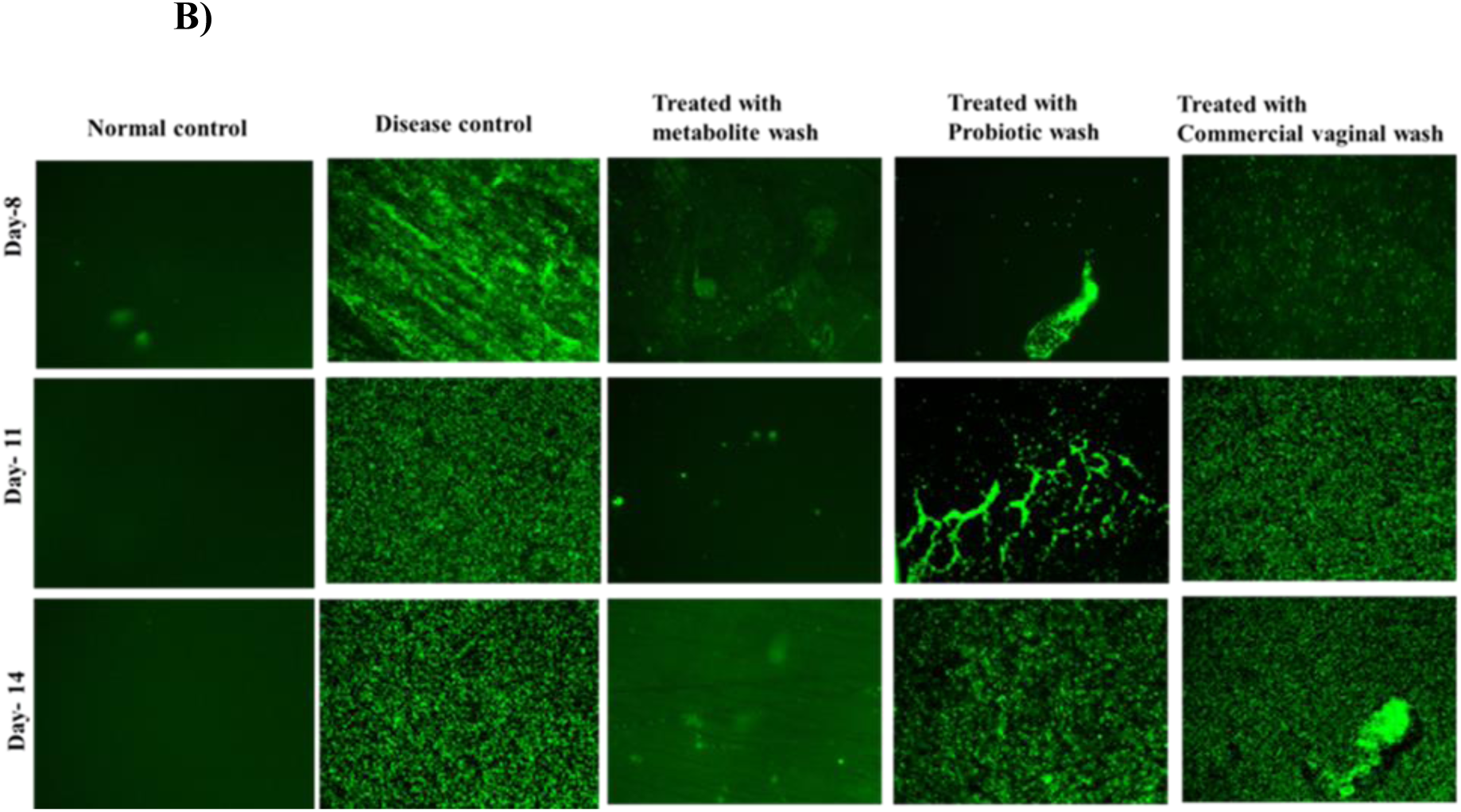
Bacterial loads in Urine of BALB/c Mice post infection. “0” on the y-axis denotes unsuccessful urine collection. Log 1 on the y-axis represents the baseline (No bacterial count) B) Fluorescent images showing the bacterial load in the urine of BALB/c Mice at Day 8, Day 11, and Day 14 post infection.

Additionally, we evaluated the bacterial burden in vital organs implicated in UTIs, including the kidney, urinary bladder, and vagina. Mice were sacrificed, and the organs were homogenized to enumerate bacterial load. Mice treated with the metabolite wash exhibited no bacterial load relative to either untreated control or treatments with probiotic wash/ commercial wash, highlighting its protective effects against cystitis and vaginal infections (Figure. 9). The normal creatinine levels were maintained in these mice during metabolite wash, further demonstrating the safety and efficacy of the metabolite wash in preventing infection and preserving renal function (Supplementary Figure. 4).Thus the formulated metabolite wash was superior in successfully treating UTI caused by UPEC in mice models relative to either a commercial wash/probiotic wash. In conjunction with non toxic nature, the customized postbiotic formulation carries a strong translational potential and can be successfully employed as a stand-alone/ supplemental therapy to treat UTI infections.

**Figure. 9.**
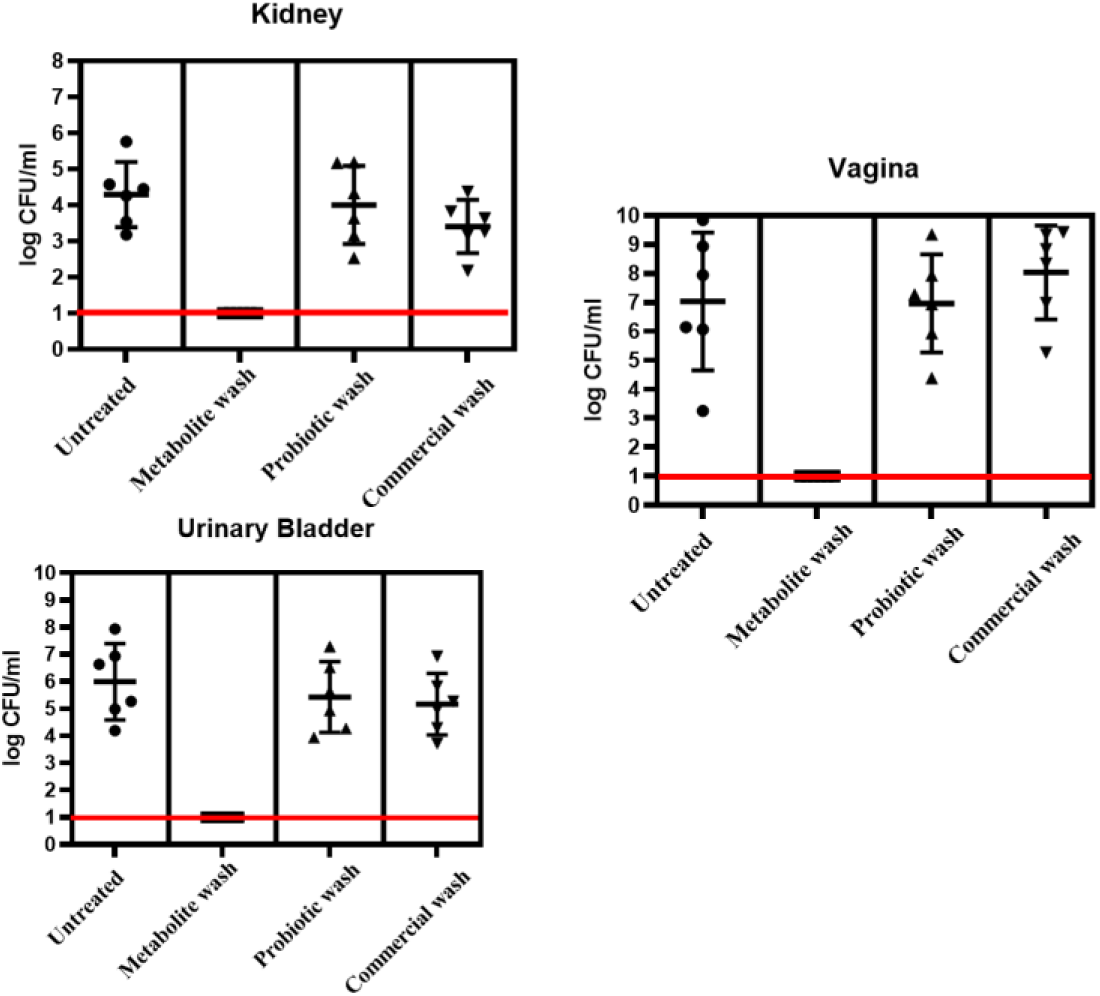
Bacterial bioburdens in organs (Kidney, Vagina and Urinary bladder) Quantifications were performed in triplicate and represented as a mean ± standard error of the mean (SEM) (N= 6)

## Discussion

Although postbiotic supplements are not as widely available as probiotics yet, postbiotics are still superior due to their purity, ease of preparation, long shelf life, capacity for mass production, precise action, and ability to elicit more targeted responses through ligand-receptor interactions.(26). Because postbiotics don’t include any bacteria—dead, live, or fragmented— they have a higher safety profile than probiotics. The increasing interest in postbiotics as a safer and more effective alternative to probiotics in feminine hygiene products is noteworthy. Postbiotics, which are non-viable microbial metabolites, offer several advantages over traditional probiotics, particularly in formulations aimed at preventing urinary tract infections Postbiotics are both natural and offer a unique and promising method for reestablishing host-microbe balance with low toxicity and consistent functional features since their lack of bacterial components reduces the possibility of potentially hazardous bacterial dissemination (Figure. 1 &Supplementary figure 1). The advantages of using postbiotics in vaginal washes are they do not contain live bacteria, which significantly reduces the risk of infections, especially in vulnerable populations. This absence of viable microorganisms means there’s no potential for bacterial translocation, making them safer than probiotic formulations that may include live or dead bacteria. Postbiotics can elicit specific responses through ligand-receptor interactions, allowing for more precise therapeutic effects. This targeted action can enhance their effectiveness in managing bacterial infections and maintaining vaginal health. The formulation of postbiotics can be more economical than probiotics, making them accessible to a broader range of consumers across different socioeconomic groups (27, 28) Due to their non-viable nature, postbiotics are easier to formulate and have a longer shelf life compared to probiotics(29). While postbiotic formulations are emerging in the market, many existing feminine hygiene products still rely on probiotics. Most probiotic-based hygiene washes incorporate live bacteria such as Lactobacillus species, which are believed to help restore the natural flora of the vagina and prevent infections. However, these products may carry risks associated with bacterial viability and potential contamination(30). The present invention relates to a novel feminine hygiene wash formulation designed to improve women’s health and well-being. The formulation comprises a combination of postbiotics and poloxamer 407, offering a fragrance-free, pH balanced solution that effectively prevents urinary tract infections (UTIs). This invention provides a cost-effective alternative to antibiotics and aims to reach all socioeconomic groups, thereby enhancing the quality of life for women worldwide.

The in *vivo* findings suggest that the customized vaginal wash offers a promising therapeutic strategy for managing bacterial infections and preventing UTIs, outperforming both probiotic and commercial formulations. Further investigation in clinical settings is warranted to validate these results and explore the potential for broader applications in human health.

## Acknowledgement

The authors acknowledge SASTRA Deemed University for the infrastructural support. We would like to express our gratitude to the Centre for Advanced Research in Indian Systems of Medicine (CARISM), supported by the FIST Biochemical Engineering-II grant (SR/FST/ET-1/2020/614), for their generous support. Additionally, we extend our sincere thanks to Ms. Sudha V for her indispensable assistance, notably in conducting GC-MS analysis. The authors would like to thank Trichy SRM Medical college and Research Centre for the collection and isolation vaginal probiotics.

