## Supplementary file for "Customized feminine hygiene wash containing postbiotics from *Lactobacillus* spp. to treat Urinary Tract Infections (UTI)"

**[b] Central Animal House Facility, SASTRA Deemed to be University, Thanjavur, 613401, India**

**[c] Pharmaceutical Technology Laboratory, School of Chemical and Biotechnology, SASTRA Deemed to be University, Thanjavur, 613401, India**

**[d] Department of Chemistry, School of Chemical and Biotechnology, SASTRA Deemed University, Thanjavur – 613 401, Tamil Nadu, India.**

**[e] Assistant Professor, Department of Obstetrics and Gynaecology, TSRMMCH&RC, Tiruchirappalli, Tamil Nadu, India**

**[f] Associate Professor, Department of Microbiology, TSRMMCH&RC, Tiruchirappalli, Tamil Nadu, India**

**[g] Research Faculty, Institutional Research Board TSRMMCH&RC, Tiruchirappalli, Tamil Nadu, India**

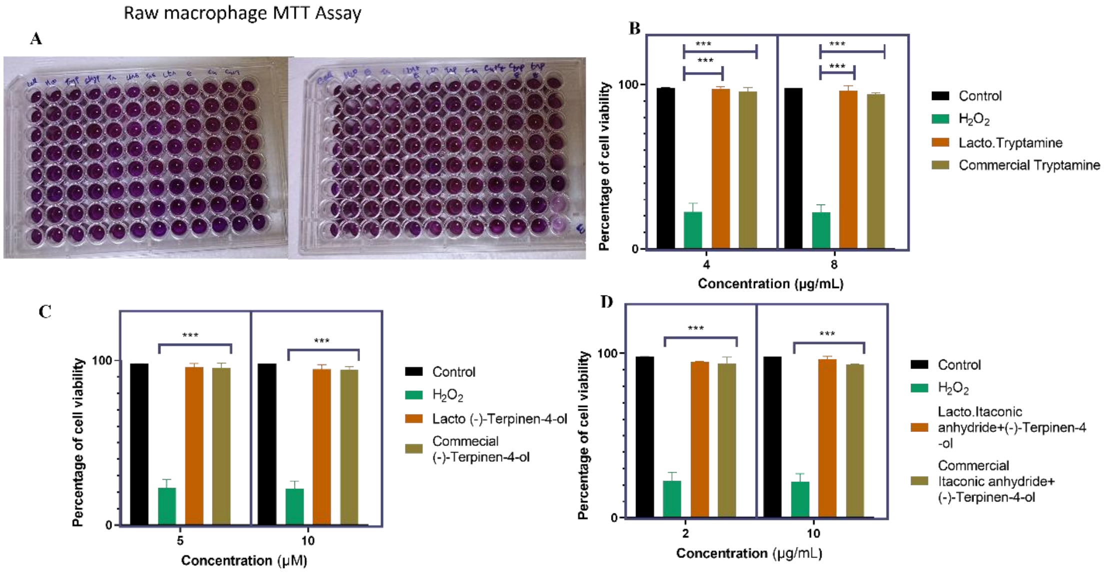


**Supplementary Figure 1.** ***Invitro*** **cell line toxicity studies for using Raw macrophages.** A) Plate showing MTT Assay B) For Tryptamine (4 µg/ml) C) (-)-terpinen-4-ol (5 µg/ml). D) itaconic anhydride (8µg/ml) $p < 0.05$ significative difference by ANOVA followed by Tukey's test.

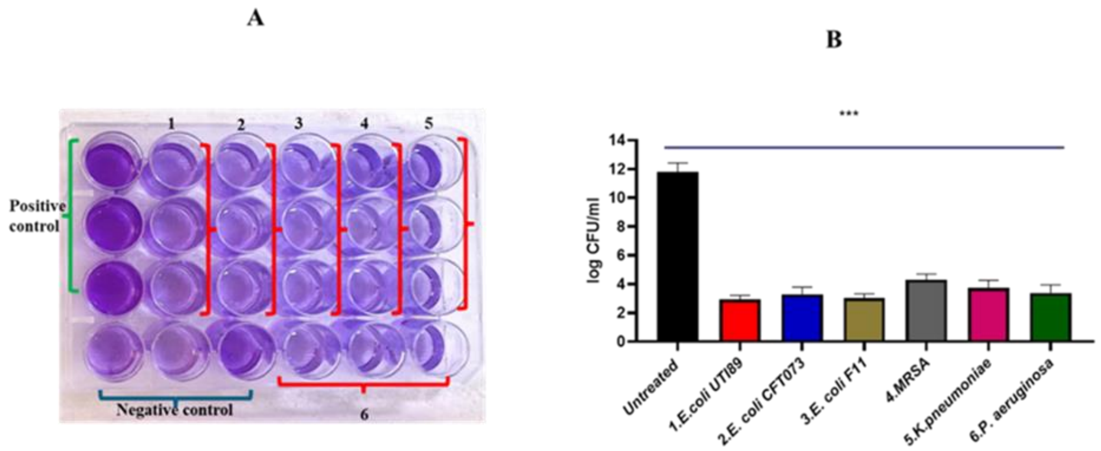


**Supplementary Figure 2. Stability Assessment of Customized *Lactobacillus*-Derived Metabolite Wash**. Crystal Violet (CV) assay (A) and Colony Forming Unit (B) count following the evaluation of the customized lactobacillus-derived wash against multiple clinical strains of bacteria. It includes wells for the Positive Control, indicating bacterial growth and biofilm formation in the absence of treatment, and the Negative Control, showing minimal bacterial growth and biofilm formation without exposure to the wash. The wells are labeled with specific clinical strains: 1. *E. coli* UTI89, 2. *E. coli* CFT073, 3. *E. coli* F11, 4. Methicillin-Resistant *Staphylococcus aureus* (MRSA), 5. *Klebsiella pneumoniae* and 6. *Pseudomonas aeruginosa*. One tailed *t* test was performed to determine the significance ***$p<0.001$ (n=3)

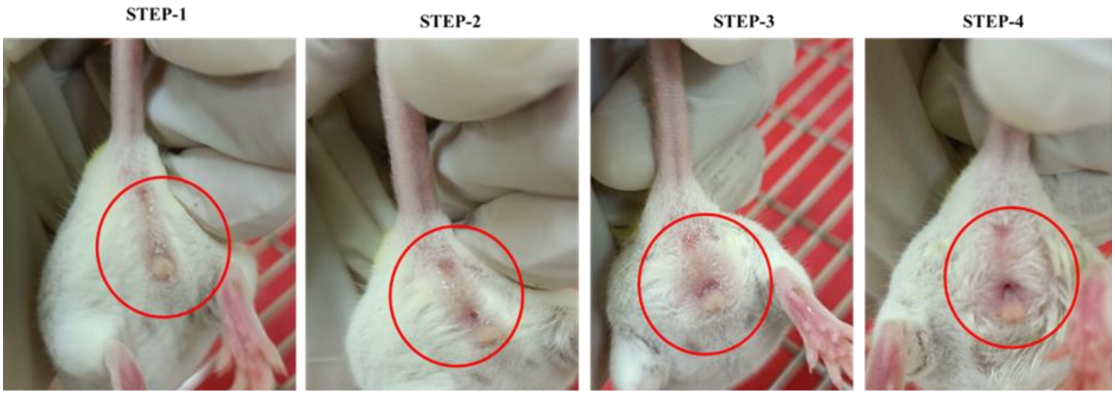


**Supplementary Figure 3: Steps showing the washing of vagina of BALB/c Mice with the formulation**. Step 1: A precise volume of 1 ml of the formulation is carefully poured from the perianal area, allowing it to flow into the vaginal opening. Step 2: The mouse is held in this position for two minutes to ensure adequate exposure to the formulation. Step 3: During this period, the formulation transitions into a gel-like consistency, which enhances its adherence and effectiveness. Step 4: After the two-minute wait, a gentle water wash is performed. A wash bottle is used to deliver the water, ensuring a thorough but gentle wash to remove the gel.

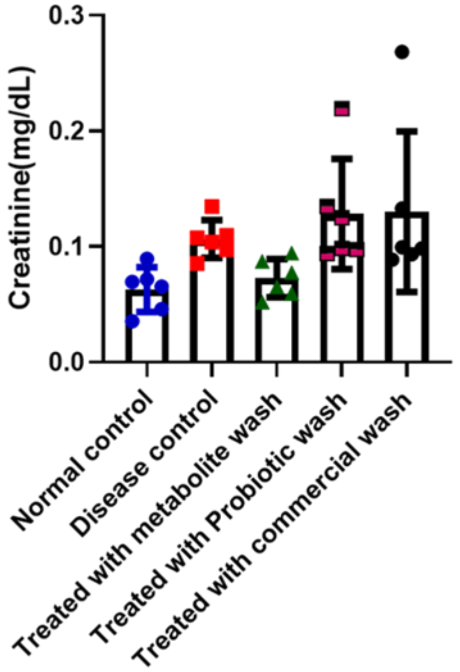


**Supplementary Figure.4: Serum creatinine Assay.** Normal serum creatinine values in BALB/c mice range from 0.08-0.11 mg/dL. Two samples got hemolyzed during blood collection time Data represent the mean ± SD.
